# Limb Position Effect in Myoelectric Control: Strategies for Optimisation and Standardisation

**DOI:** 10.1101/2025.09.01.673545

**Authors:** Trevor Overton, Zubaidah Al-Mashhadani, Syed Yahya Raza, Jason Whitson, Mohsen Rakhshan

## Abstract

**Objective:** Myoelectric control uses electromyography (EMG) signals for muscle-machine interfacing with applications in prostheses, augmented/virtual reality, and consumer electronics. However, factors such as changes in the limb’s position during activities of daily living reduce the controller’s reliability. Therefore, there is a need to develop techniques that reduce this limb position effect to increase the widespread adoption of these technologies.

**Approach:** We developed an open-source device to standardise myoelectric control experiments. The device has sixteen locations for automatically positioning the participant’s arms to perform hand gestures or grasp objects, with lights and sensors for guidance and timekeeping. We used this device to collect data from eighteen healthy participants in a five-session study under three modalities: performing five hand gestures with a static or dynamic limb and moving three objects. We recorded forearm electromyography and kinematics of the upper limb and trained a linear discriminant analysis model to assess the classifier’s accuracy across different modalities and arm positions.

**Main results:** The classifier’s accuracy with a static limb was decreased when tested on untrained positions, confirming the limb position effect. More training positions improved accuracy, with four optimally balancing the training burden and classifier accuracy. Classifiers trained with data from dynamic movements outperformed when tested on dynamic data. Furthermore, adding kinematic data to the classifier increased accuracy yet significantly reduced learning rates. However, training with a dynamic limb improved this learning rate.

**Significance:** The limb position effect can be countered by training with multiple positions and including kinematic data. Classifiers with EMG and kinematic data should be trained using a dynamic limb to achieve high accuracy with reasonable amounts of training data. Our open-source, automated device will help standardise datasets between laboratories, aiding the further development of robust and widespread myoelectric control.

## 1. Introduction

Myoelectric control uses electromyography (EMG) signals and machine learning techniques to manipulate a robot or computer [1, 2]. Decades of research have resulted in advanced systems that can accurately determine muscle activation patterns [3-9]. Numerous applications arise from this technology, such as control of upper-limb prostheses [3, 7-11], wheelchairs [12, 13], speech decoding devices [14, 15], augmented and virtual reality (AR/VR) [16, 17], and consumer electronics [18, 19]. These applications require robust and accurate decoding of movements to create the best user experience.

Despite high decoding accuracies within laboratory-controlled conditions, myoelectric control is known to have several confounding factors that reduce its usability during activities of daily living (ADL). These factors include the limb’s position, fatigue, contraction intensity, electrode shift, and across-day effects [20-22]. In prosthesis control, such confounding factors result in an unreliable controller and have been cited as a major reason for abandoning the prosthesis [23]. Therefore, research must be performed to help bridge the gap and create functional control during ADL to help increase the model’s robustness to the confounding factors.

One major confounding factor that impacts the accuracy of myoelectric controllers is the limb position effect. More specifically, the limb position effect describes how changes in the limb position, compared to the position in which the classifier was trained, increase the error of the classifier [24]. Some methods, like training the classifier in multiple positions, have been shown to reduce this effect [24-27]. However, training on multiple positions increases the training burden, making it less desirable if the myoelectric controller needs to be re-trained for a long time before every use.

Alternative approaches for countering the limb position effect have also been researched. For instance, reducing motor variability by training participants in a closed loop, similar to neurofeedback, is found to be effective [28, 29]. In addition, numerous studies have shown promise in a multi-modal approach of inputting inertial and EMG data into classifiers [24, 30-32], but the classifiers require extra training information from many positions to achieve improved robustness [30]. Therefore, these alternative approaches often fail to reduce the training burden, which is essential for the widespread adoption of myoelectric control.

In the past few years, attention has turned to reducing the training burden by creating algorithms for devices that could be used “off-the-shelf”. Deep-learning neural networks have already shown some resilience to the limb position effect [33]. Further, Meta has recently proposed that such a deep learning network is effective across people, with no training data required from new users [34]. However, their models are challenging to embed into a small microcontroller with limited power.

Beyond the limb position effect and training burden, the limbs are also continuously moving during ADL. Therefore, it is essential to study how a dynamically moving limb could affect myoelectric control. Previous studies have found that including training data with dynamic movements was crucial to reducing errors and increasing robustness [35-38]. Furthermore, it is shown that using dynamic movements could help improve the learning rate of a classifier when EMG and inertial data are combined [30]. A recent study even showed that the choice of electrode position can impact the robustness of classifiers under dynamic limb conditions [39].

Our study focuses on optimising myoelectric control for robustness to the limb position effect. We develop an open-source device to standardise the assessment of the limb position effect across both static and dynamic limb conditions, and use it to collect a large dataset with minimal experimenter’s interference. We demonstrate strategies to overcome the limb condition effect, with minimal training burden, for optimal myoelectric control. Importantly, these results should be reproducible in the community with our open-sourced device, setting a standard for future studies to build upon.

## 2. Methods

This study was performed in accordance with the Declaration of Helsinki. This human study was approved by the University of Central Florida’s Institutional Review Board (study 6576). All adult participants provided informed consent to participate in this study.

To study the limb position effect of myoelectric control, we developed an open-source device to collect data from participants. All details to replicate the device are placed on the lab’s GitHub page (https://github.com/LIMB-UCF/CAPSAS_Documentation). Due to its documented and open-source layout, this low-cost device can be used to standardise studies seeking to develop robust myoelectric control for muscle-machine interfaces.

The Common Arm Position Signal Acquisition System (CAPSAS) device spans sixteen distinct arm positions (figure 1). Each arm position is outlined by a hexagon that contains twelve LEDs and a central infrared (IR) proximity sensor. Using the LEDs, specific animations are presented to the participants to instruct them on what hand gesture to perform. When the arm is moved to the location, the IR proximity sensors timestamp the movement automatically. With its automated nature, the CAPSAS removes the experimenter’s interference and streamlines the data collection process.

**Figure 1.**
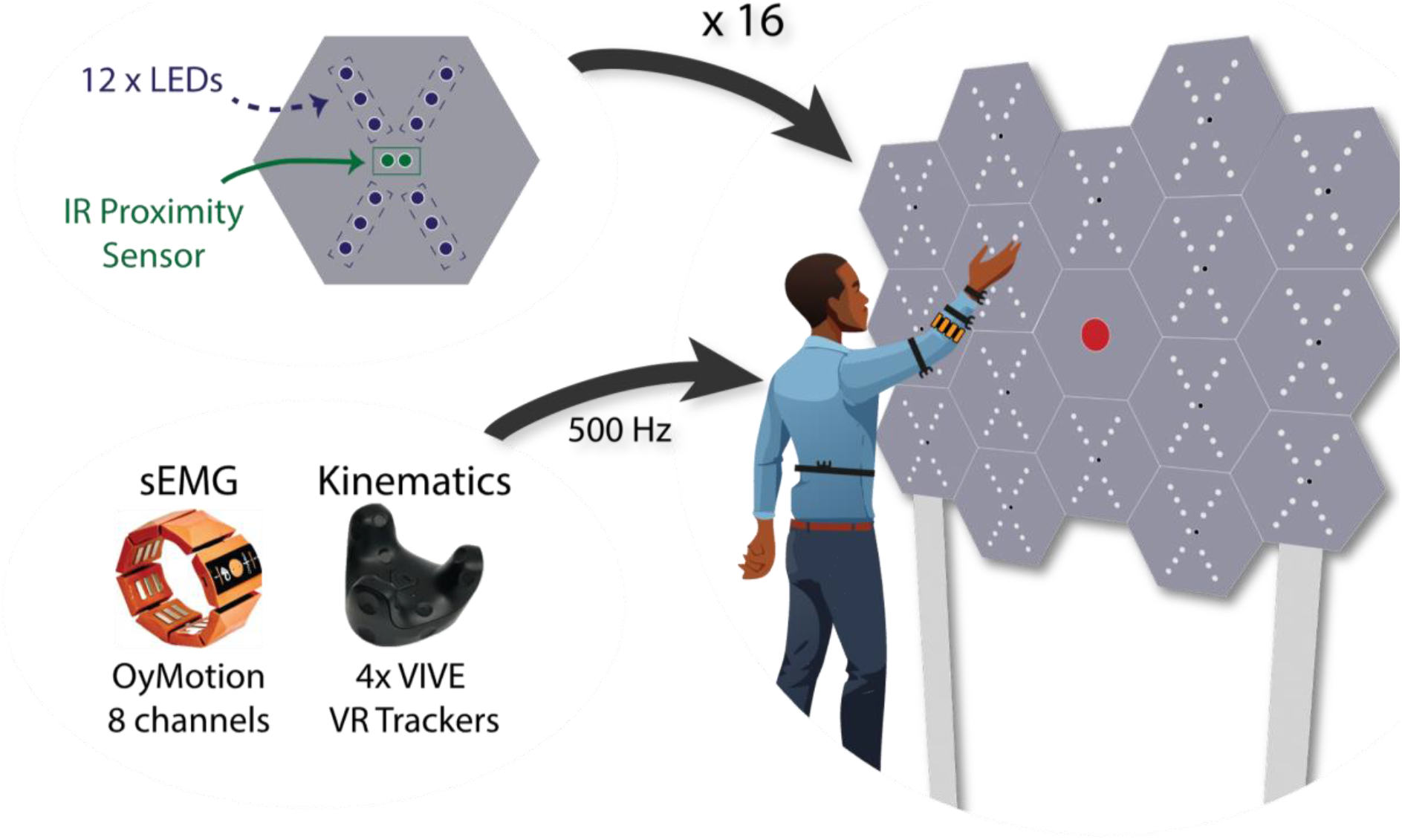
Common Arm Position Signal Acquisition System (CAPSAS) comprises sixteen arm positions, each with its own proximity sensor and LEDs. The lights guide a participant in performing hand gestures in front of the device. At the same time, a sEMG armband (OyMotion gForcePro+) records eight channels of muscle activity and four VR motion trackers (HTC VIVE tracker 3.0) record kinematics.

While the participants were directed by the CAPSAS, eight surface EMG (sEMG) channels were recorded from the forearm using an OyMotion gForcePro+ armband. The kinematics of the upper limb were tracked using four HTC Vive motion trackers. All data was recorded at 500 Hz and synchronised using the Lab Streaming Layer Python library [40].

Eighteen healthy participants were recruited for the study (fourteen male and four female; fifteen right-hand dominant). Every participant took part in a total of five sessions spanning across days, with each session lasting around 75 minutes. Participants performed in three modalities: static, dynamic, and object modality (see below; static and dynamic lasted two sessions each). The different modalities allowed us to study various aspects of the limb position effect of myoelectric control. The order of the modality and the trials within each session were counterbalanced to prevent any order effect.

Due to Bluetooth connectivity issues during a few sessions, we decided to exclude the data from six sessions to maintain data integrity. More specifically, two static sessions (from participants 8 and 18), two dynamic sessions (from participants 7 and 9), and two object sessions (from participants 4 and 5) were removed from the data analyses.

### 2.1 Static Modality

In the static modality, participants performed five hand gestures at all sixteen positions: hand open, hand closed, wrist extension, wrist flexion, and rest (figure 2(a)). The participant performed each hand gesture for five seconds while holding their hand in front of the corresponding position on the CAPSAS.

**Figure 2.**
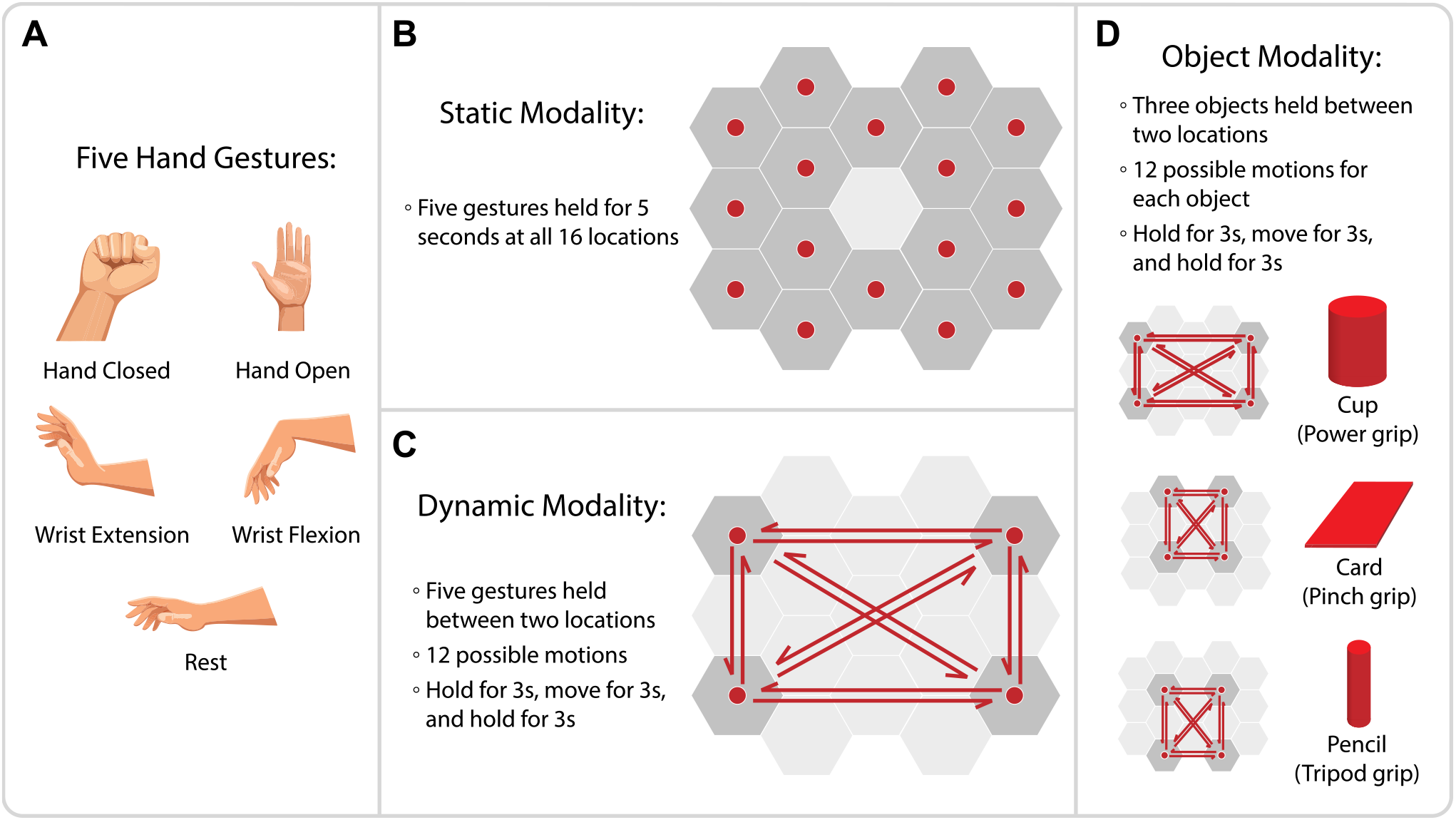
Data collection methods. (a) Five hand gestures were used in the static and dynamic modalities: open, closed, extension, flexion, and rest. (b) The static modality had participants hold each of the five gestures at all sixteen locations while holding their limb still for five seconds. (c) The dynamic modality had participants hold each of the five gestures while moving their arm. Each trial consisted of three seconds at one location, a three-second motion to another location, and three seconds at the final location. Twelve possible motions spanning the corners of the device were tested. (d) The object modality had participants move three objects (cup, card, and pencil) through twelve possible motions, timing similar to the dynamic modality.

Each participant took part in two sessions of the static modality across two different days. Each session required the participant to perform five gestures at all sixteen positions three times, for a total of 240 trials. Every 80 trials (i.e., one block), there was a five-minute break to prevent fatigue. Notably, the automated design of the CAPSAS allowed the participant to proceed through the trials in each block at their own pace, with no limit on the inter-trial rest period.

### 2.2 Dynamic Modality

The dynamic modality used the same five gestures as the static one; however, the participant moved their limb in a structured path while performing the gesture. More specifically, they started at one position for three seconds, moved to another for three seconds, and finally held the gesture in that position for the last three seconds (figure 2(c)). The participants were instructed to follow a straight path between the two positions. Notably, the LEDs on CAPSAS instructed the participants of the starting and ending positions. Furthermore, a dynamic colour change occurred in all the positions except the starting and ending positions, informing the participant about the remaining time of the trial. Before the data collection, the participants performed a familiarisation task to get used to the experiment and the colour’s meaning. Since testing all possible movements would be highly time-consuming, twelve different movements spanning the device’s corners were used to study the maximal effects of the limb’s motion (figure 2(c)).

The dynamic modality was structured so that the static segments (beginning and ending three seconds) could be compared to the dynamic segment (middle three seconds) by splitting the data with time windows based on the IR proximity sensors. Therefore, we can thoroughly investigate how myoelectric control is affected by a moving limb, which is typical during ADL. We should note that the second static segment suffered a major deterioration in quality compared to the other segments, as measured by its classification accuracy (supplementary figure 1). The effect may be due to participants trying to activate the CAPSAS IR sensor at the end location by moving the arm and hand. Therefore, our analyses only considered the first static segment and dynamic segment, ensuring any effects found are most likely due to the limb motion.

Each participant took part in two sessions of the dynamic modality. Each session required the participant to perform five gestures through all twelve motions three times for a total of 180 trials. Every 60 trials, there was a five-minute break to prevent fatigue. Similar to the static modality, the participants were allowed to proceed through the trials at their own pace, with no limit on the inter-trial rest period.

### 2.3 Object Modality

The object modality comprised twelve dynamic motions while the participant grasped three objects (3D-printed cup, card, and pencil). The CAPSAS device has customisable magnet object holders, facilitating experimentation with real-life objects. Due to the physical positioning of the objects on the CAPSAS device, each one had a slightly different path that was used (figure 2(d)). Notably, these differences in paths due to object class led us not to use the motion tracker data to analyse this modality, as they would contain the information about the object.

Like the dynamic modality, a nine-second pattern (three seconds at the first position, a three-second movement, and three seconds at the final position) was used for each trial. Since the object modality was structured similarly to the dynamic modality, investigations into the effect of motion on myoelectric control could be conducted. Furthermore, using objects allowed for studying myoelectric control independent of discrete gestures, which are unrealistic for some myoelectric control applications in daily life or advanced AR/VR environments.

Each participant took part in one session of the object modality. Each session required the participant to move the three objects through all twelve motions six times for a total of 216 trials. Every 72 trials, there was a five-minute break to prevent fatigue. Similar to the other modalities, participants were allowed to proceed through the trials at their own pace, with no limit on the inter-trial rest period.

### 2.4 Data Processing and Classification

Every trial was segmented into time windows of 100 ms (50 samples) with a step size of 100 ms (50 samples). The step size was chosen to prevent overlap in the training and testing data of the classifier.

A linear discriminant analysis (LDA) model was used to classify the hand gestures using mean absolute value, variance, zero crossings, slope change, and waveform length of the EMG data within the window [6, 41, 42]. The mean absolute value of the EMG data within a time window was defined as

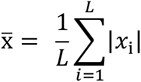

where *L*is the length of the window and *x*_*i*_ is the *i*th sample in the window. The variance of the EMG data within a window was defined as

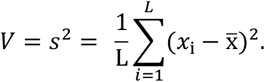

Zero crossings were counted every time the waveform crossed zero, with a threshold of ε at 2 µV [41]. The count was increased for each two consecutive samples, *x*_*i*_ and *x*_*i*+1_, such that

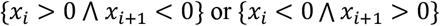

and

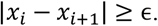

Slope sign changes were counted by first finding the gradient of the EMG data within a time window and then counting the zero crossings with a similar threshold of *ϵ* (2 µV). Finally, the waveform length was defined as

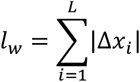

where Δ*x*_*i*_ = *x*_*i*+1_ − *x*_*i*_.

When used for the analyses, the kinematic data was included as averaged quaternions as additional features for the classifier. Each tracker had four quaternion values (*x, y, z, w*) and the mean of the value over the time window was included as features (four features per motion tracker).

### 2.5 Analysis of the Classifier’s Performance and Learning Rates

The classifier was tested on unseen data, and its percent accuracy was recorded. Several parameters were used to compare the classifiers’ performance to each other. When necessary to find the optimal point considering a trade-off between the amount of training data and accuracy, we calculated the knee point of the learning curves [43].

The Shapiro-Wilk test was used to assess the data’s normality. Since the accuracies measured were not normal, we used non-parametric hypothesis tests. Further, to analyse the effect size, the *r* value was calculated by taking the *z* statistic of the test and dividing by the square root of the sample size.

Several of the analyses examined the learning rate of the classifier. In these analyses, the classifier was trained on an increasing number of trials from the session, and its accuracy was tested on the remaining trials. The resulting curve of training data versus classifier accuracy was fit to a hyperbolic curve

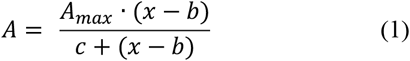

where *A* is the accuracy of the classifier, *A*_*max*_ is the horizontal asymptote, and τ = *b* + *c* is the number of trials required to reach an accuracy of 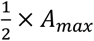. The *A*_*max*_value should be constrained between zero and one in the curve fitting algorithm, since an accuracy greater than one is impossible. The overall accuracy of the models could be compared with their *A*_*max*_ values and the learning rates through their τ values.

## 3. Results

### 3.1 Confirmation of the Limb Position Effect

Myoelectric control needs to be consistent and reliable during activities of daily living (ADL) or when interacting with virtual or augmented reality (VR/AR) environments. However, many confounding factors limit this reliability. One important factor is the position of the limb (i.e., the limb position effect), in which the limb’s position can deteriorate the accuracy of myoelectric control. To standardise the examination of this effect, we developed an open-source device called the Common Arm Position Signal Acquisition System (CAPSAS). The device spans sixteen distinct arm positions and enables the researchers to study the limb position effect in different settings, such as performing hand gestures with static or dynamic limbs or moving real-life objects in a controlled and automated fashion with no experimenter’s interference (figure 1; see Methods section for details). This device enabled us to examine the limb position in different settings, as explained below.

Eighteen healthy participants performed in five sessions with three modalities: static, dynamic, and object. In the static modality, which was collected across two sessions, participants performed five hand gestures (open, close, wrist extension, wrist flexion, and rest; figure 2(a)) for five seconds in front of all sixteen positions on CAPSAS.

First, we wanted to confirm the existence of the limb position effect in the data collected using CAPSAS in both sessions of the static modality. Therefore, we compared the classification accuracy when the classifier was trained and tested on the same position (i.e., intra-position accuracy) with when the classifier was trained on one position and tested on all other fifteen positions (i.e., inter-position accuracy).

Specifically, we trained a linear discriminant analysis (LDA) classifier with mean absolute value, variance, zero crossings, slope change, and waveform length of the EMG data within the window as the input features (see Methods section for details). We found that the classifier’s accuracy in the inter-position setting was significantly lower than in the intra-position setting (81.6 ± 12.8% vs. 92.3 ± 6.3%, *p* ≈ 0, *r* = 19.875, Wilcoxon signed-rank; figure 3(a)). Therefore, the limb position effect was present in our collected data.

**Figure 3.**
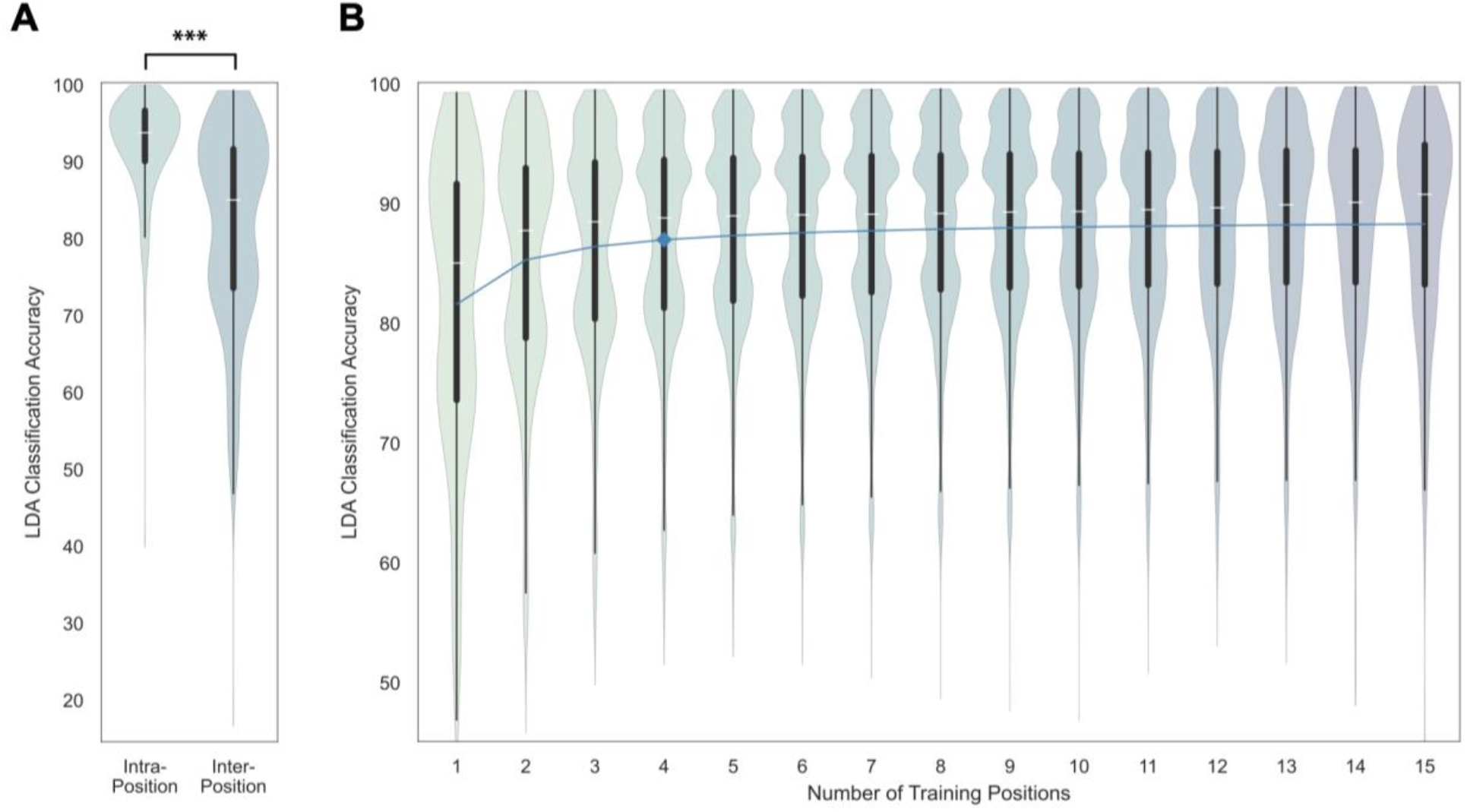
Analysis of the static modality shows the limb position effect. (a) LDA models were trained in one arm position and tested either within the same position (intra-position) or between different positions (inter-position). The intra-position accuracy was significantly higher than the inter-position accuracy (Wilcoxon signed-rank, *p* ≈ 0, *r* = 19.875). (b) LDA models were trained on an increasing number of arm positions and then tested on all the remaining positions. The blue line tracks the mean accuracy, which increases with every training position added. The number of positions optimally balancing the increasing training burden and accuracy is marked at four.

#### 3.1.1 Using Four Arm Training Positions Is Ideal for Robustness to the Limb Position Effect

Observing the limb position effect, we next explored the strategies that can reduce this effect. Research has shown that increasing the arm positions on which the classifier is trained can increase its reliability [24]. However, one drawback associated with this approach is an increased training burden. Hence, an ideal number of arm positions should be determined to optimise the trade-off between quick training data collection and robustness to limb position effect. Therefore, we used the data from the static modality to train the classifier on a subset of the positions and test on the remaining positions. We found that by increasing the number of positions used for training the classifier, the testing accuracy indeed improved (figure 3(b)). However, the rate of increase diminished as more positions were added to the training data. We used the knee point of the accuracy curve to find the optimal number of positions that balance the training burden and accuracy (see Methods section for details). This analysis suggested that training the classifier on four positions results in an optimal trade-off between quick training data collection and robustness to limb position effect.

### 3.2 Documenting the Effects of Dynamic Limb Motion

The limb position effect is often studied in the context of a static limb in different positions. However, during activities of daily living or when interacting with AR/VR environments, the limbs constantly move, and the intended hand gesture needs to be reliably decoded throughout the movement.

Therefore, we designed the dynamic modality experiment and collected the data using CAPSAS. In this modality, which was performed over two sessions, the participants were instructed by the device to move their arm to a specific position and perform a particular hand gesture for three seconds without moving the arm. Then, the device instructed the participant to move their arm to another position. Finally, the gesture had to be kept at the ending position for three more seconds. The design allowed us to compare how the classifier accuracy is affected by the active movement (i.e., motion) of the arm.

#### 3.2.1 Limb Motion Does Not Significantly Impact Classifier Accuracy

Using the dynamic modality, we first tested whether the motion of the limb negatively impacts the classifier accuracy. To examine this question, we split the data of each trial within the dynamic modality into three segments based on the motion of the limb. The first three seconds in which the arm was held in front of the starting position will correspond to the first static segment. The middle three seconds of the trial, in which the arm moved from the starting position to the end position, were segmented and called the dynamic segment. Finally, the last three seconds, in which the hand was held in front of the end position, were segmented and called the second static segment.

To remove any motion artefacts from the two static segments in the following analysis, we only included a one-and-a-half-second window, which was one second after moving to the intended position and half a second before the end of the segment. Furthermore, to ensure the data in the dynamic segment had motion, we excluded the first half-second of data from when CAPSAS signalled the participant to move. To ensure the number of data points between each static segment and the dynamic segment is equal, we also used a one-and-a-half-second window for the dynamic segment. Additionally, to examine how the classifier performs across both dynamic and static segments together, a special window of data was used. In this window, the data started from one second after the trial began and lasted for four seconds. Therefore, the segment contained two seconds of static data and two seconds of dynamic data.

We trained the classifiers on the first static segment of all positions in the dynamic modality (see Methods for exclusion of the second static segment). We tested on either unseen first static segment data or the dynamic segment. Two-thirds of the data (120 trials) was used to train the classifiers, while the remaining data was used to test them. We found that the classifier that was trained on the static segment did not show a significant accuracy difference when tested on the static segment versus when tested on either both segments (*p* = 0.054; Wilcoxon signed-rank test) or the dynamic segment (*p* = 0.071; Wilcoxon signed-rank test; figure 4(a)). The result shows that a classifier trained on the static arm positions (four positions) can be robust, even when the limb dynamically moves within the training positions. Importantly, this result holds even with various proportions of data used for training (supplementary figure 2).

**Figure 4.**
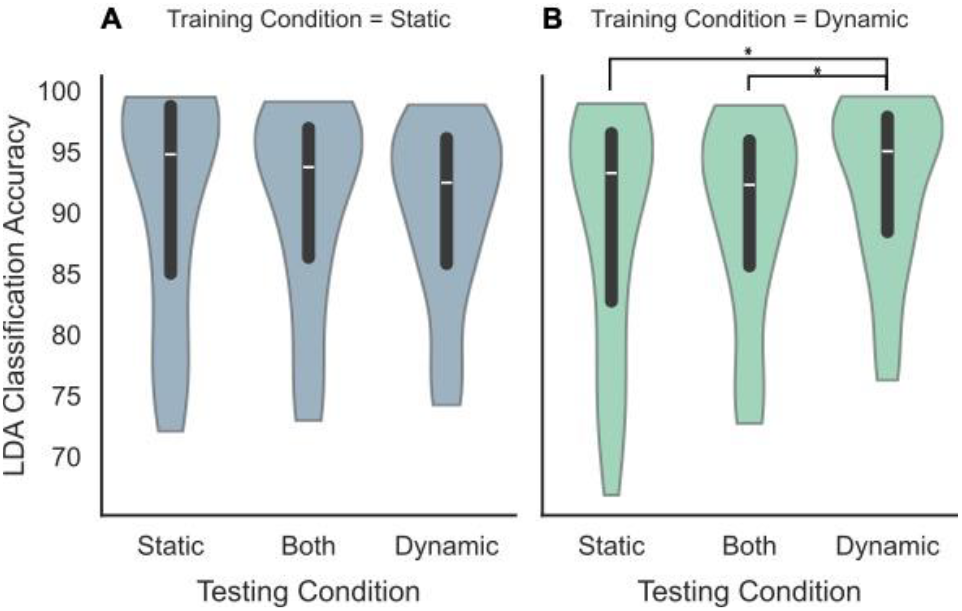
Analysis of the effects of training and testing conditions on classification accuracy during the dynamic modality. (a) LDA classifiers are trained on only the static segment of the dynamic session. Then, they are tested on either the static segment, the dynamic segment, or both segments. Two-thirds of the available data (120 trials) were used for training, and the rest for testing. Box plots of the classification accuracy are shown. A Wilcoxon signed-rank test returned no significant differences between the testing conditions. (b) Similar to (a), except the LDA classifiers are trained with the dynamic segment. A Wilcoxon signed-rank test (α = 0.05/3 = 0.017; corrected for multiple comparisons) returned two significant results. The model with dynamic testing performed better than classifiers tested with static segment data (*p* < 0.001, *r* = 0.701) and classifiers tested with both types of data (*p* < 0.001, *r* = 0.856). Additionally, a Mann-Whitney U test returned no significant differences between the dynamic and static (part (a)) training conditions.

#### 3.2.2 Classifiers Trained with Motion Outperform When Tested with Motion

We showed that training the classifier in four different arm positions is ideal for a trade-off between training burden and accuracy. However, it is unclear whether training the classifier with a moving arm will provide additional benefits. Therefore, we examined whether training the classifier on dynamic segments improved the classifier’s accuracy. More specifically, we first trained the classifier on the dynamic segment of the dynamic modality. Then, we tested the classifier on the unseen dynamic segment, static segment, and segment containing both data types (see previous section). By comparing the accuracy of these classifiers with those trained with the static segment of the prior section, we found no significant difference between training with or without motion (*p* = 0.151 between static testing, *p* = 0.448 between testing with both, and *p* = 0.128 between dynamic testing; Mann-Whitney U test; figure 4). The result suggests that training the classifier with dynamic arm motions might not provide additional benefit for classification accuracy compared to training with static data.

Notably, if the testing data included only the dynamic segment, training the classifier with the dynamic segment was indeed beneficial (figure 4(b)). More specifically, the classifiers tested with dynamic segment data outperformed both the classifiers tested with static segment data (*p* = 7.4 × 10^−5^, *r* = 0.701, Wilcoxon signed-rank) and classifiers tested with both types of data (*p* = 1.3 × 10^−6^, *r* = 0.856, Wilcoxon signed-rank). The result suggests that dynamic training could be beneficial if the prosthesis or other device is expected to be predominantly moving during control. The trend here is also consistent with various proportions of training data (supplementary figure 3).

#### 3.2.3 Object Modality Challenges Classifier Accuracy

In the analyses above, the participants performed hand gestures often used for myoelectric control, such as hand open, close, relax, and wrist flexion or extension. However, while manipulating objects in VR/AR or real life using a prosthesis, intuitive myoelectric control is essential for natural interaction. Therefore, we designed the object modality to examine how the grasp according to the shape of the object affects classification. Importantly, we did not specify the exact fingers to grasp the objects, as in the real world, there are some subtle individual differences. Each trial started with grasping an object without arm movement for three seconds (i.e., first static segment). Then, the CAPSAS device instructed the participant to transport the object to another position for three seconds (i.e., dynamic segment). Finally, the participant had to hold the object at the end position for another three seconds (i.e., the second static segment).

Interestingly, the first noticeable aspect of the object modality classification is the overall lower accuracy. The classical hand gestures used during the dynamic modality yielded median accuracies of around 95% (figure 4), while the object modality returned a median accuracy of 80% (figure 5). The object modality also had a pronounced increase in variation compared to the dynamic. The lower accuracy of the classifier in object modality, even with fewer classes, highlights a potential challenge in myoelectric control with non-classical hand gestures. This challenge remained pronounced no matter the amount of training data used (supplementary figure 4).

**Figure 5.**
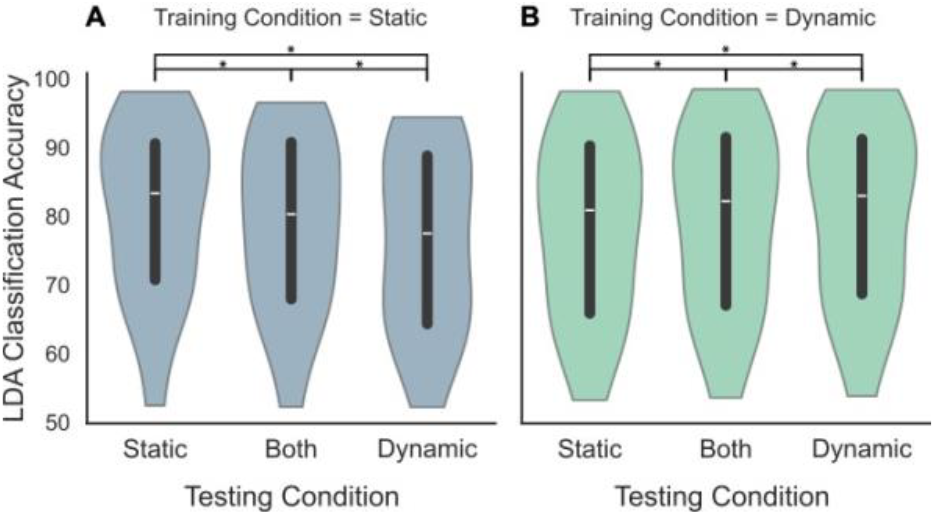
Analysis of the effects of training and testing conditions on classification accuracy during the object modality. (a) LDA classifiers are trained on only the static segment of the object session. Then, they are tested on either the static segment, dynamic segment, or both segments. For equal comparison to the analysis in figure 4, the training dataset consisted of 120 trials. Unseen data was always used for testing. Boxplots of the classification accuracy are shown, with individual sessions shown as dashed lines. A Wilcoxon signed-rank test (α = 0.05/3 = 0.017) returned significant differences between all testing conditions. (b) Similar to (a), except the LDA classifiers are trained with the dynamic segment of the object modality. A Wilcoxon signed-rank test (α = 0.05/3 = 0.017) returned significant results for all testing conditions. Additionally, a Mann-Whitney U test returned no significant differences between the dynamic and static (part (a)) training conditions.

Compared to the dynamic modality, the object modality had an even more pronounced effect when training the classifiers with data from the dynamic segment. Similar to the dynamic modality, these models saw improved performance when tested on dynamic segments (figure 5(b)). More specifically, these models tested with the dynamic segment outperformed the static segment testing (*p* = 0.002, *r* = 0.776, Wilcoxon signed-rank) and testing with both segments (*p* = 0.001, *r* = 0.621, Wilcoxon signed-rank). However, unlike the dynamic modality, the models trained with the static segment performed worse when tested with data from the dynamic segment (figure 5(a)). In particular, the models tested with the dynamic segment underperformed the static segment testing (*p* = 0.001, *r* = 0.827, Wilcoxon signed-rank) and testing with both segments (*p* = 0.001, *r* = 0.853, Wilcoxon signed-rank). Furthermore, similar to the dynamic modality, no significant differences were found between the two training conditions (*p* = 0.510 between static testing, *p* = 0.955 between testing with both, and *p* = 0.376 between dynamic testing; Mann-Whitney U test; figure 5).

### 3.3 Considerations for Kinematic Data While Minding the Training Burden

Thus far, our analyses have been limited to classification models that only have information from the EMG signals. However, we have not yet examined how much including kinematic data could affect the limb position effect. The models should also be assessed to determine whether such an inclusion will benefit the accuracy at the cost of an undesirable learning rate.

Different subsets of the four IMUs (wrist, forearm, arm, and trunk) were analysed to determine how much information is needed for an optimal balance of performance and minimal training data. The first four subsets considered consisted of each individual tracker being appended to the classifier’s features (see Methods section for details). Additionally, a subset that contained all the trackers besides the wrist was used, as it could mirror a myoelectric control application in rehabilitation engineering. Furthermore, this subset was useful as the wrist IMU could potentially hold class information, as its orientation during the hand gestures could be class-specific. However, myoelectric control contexts besides rehabilitation could benefit from this additional information. Hence, a set of all IMUs was also analysed.

We trained classifiers with the various subsets of IMU data across different numbers of trials for training data (figure 6). Then, we created a learning curve of the models by plotting accuracy as a function of the number of trials for the training data. The learning curves allow for an analysis of both classification accuracy and learning rates (see below).

**Figure 6.**
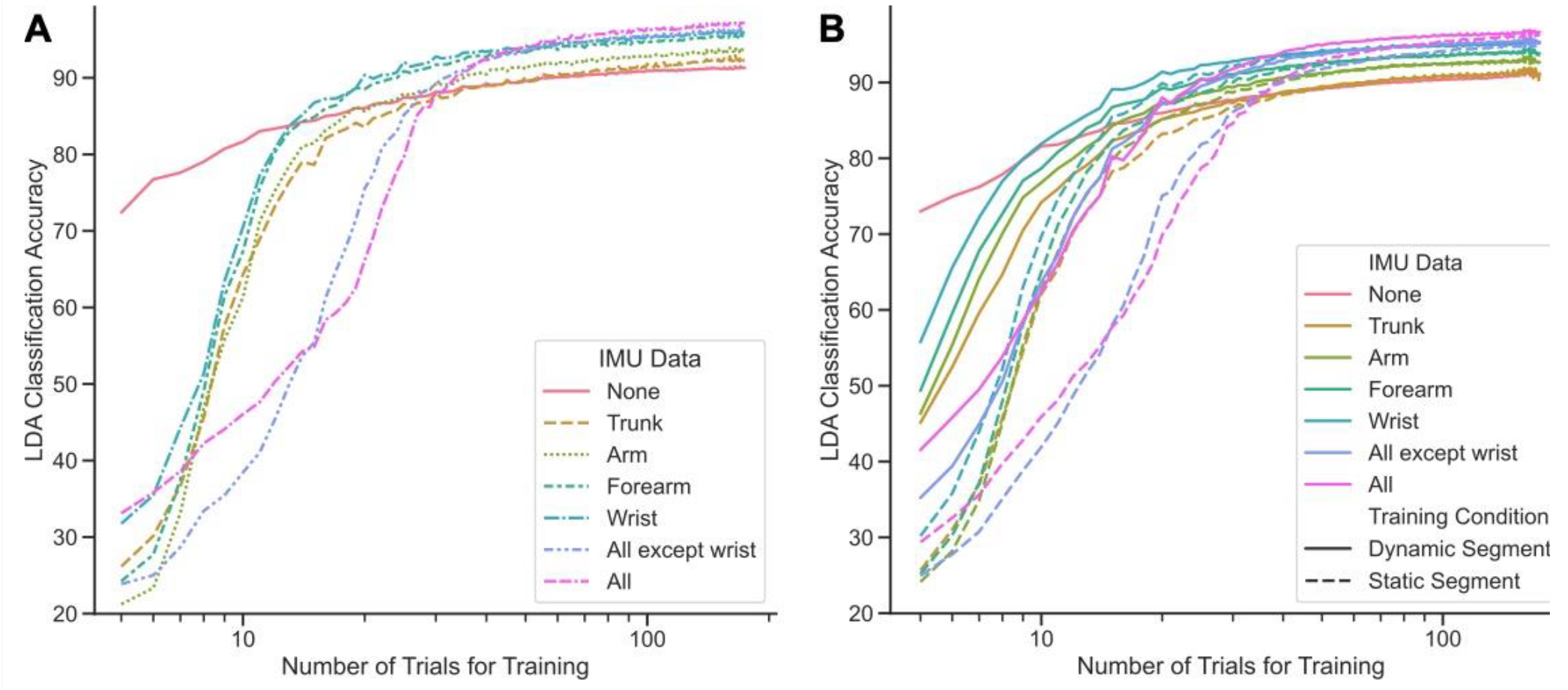
Analysis of the classifier’s learning curves with the inclusion of kinematic data. (a) Comparison of different IMU data inclusion during the static modality. The solid line shows the learning curve of LDA models trained without IMU data. Various dashed lines show different subsets of IMUs being included in the model. (b) Comparison of different IMU data inclusion during the dynamic modality. Various colours depict the different subsets of IMUs being included in the model. Solid lines show models trained from a dataset of the dynamic arm motions, while the dashed lines show models trained from a dataset where the arm was static. The testing dataset consisted of both portions of static and dynamic segments.

#### 3.3.1 Kinematic Data Increases Accuracy Yet Decreases Learning Rates

Overall, in the static modality, the IMU data provide increased classification accuracy with large training datasets (figure 6(a)). More specifically, the *A*_*max*_ parameter from equation (1) can be used to compare the hypothetical accuracy of the classifiers with infinite training data. While the model with no IMU data has an average accuracy of 92.7% ± 8.8%, all the models incorporating IMU data performed significantly better. The individual models can be listed by the order of increasing performance: trunk IMU (94.4% ± 6.4%), arm IMU (95.5% ± 5.1%), forearm IMU (97.1% ± 3.4%), and wrist IMU (97.5% ± 3.7%). Additionally, the multiple IMU models performed even better, with average accuracies of 99.4% ± 1.5% when all IMUs except the wrist are included and 99.7% ± 1.0% when all IMUs are included.

Despite the benefits, these classifiers displayed a slower learning rate than a model that had no kinematic data (figure 6(a)). To quantify the impact of including kinematic data on the learning rate, we fit each participant’s data to equation (1) and compared the parameter τ. We found that the classifiers trained without kinematic data had an average value of 2.95 ± 1.2. However, all classifiers with one IMU have average τ values greater than six (trunk: 7.3 ± 1.9; arm: 7.4 ± 1.5; forearm: 6.9 ± 1.4; and wrist: 6.5 ± 0.8). Furthermore, the classifiers with multiple trackers’ data included had an average τ of greater than nine (all except wrist: 10.0 ± 2.5; and all: 9.4 ± 2.3). These results show that including more kinematic data means the classifiers learn at a slower rate, resulting in a greater training burden.

#### 3.3.2 Training with Dynamic Arm Motions Counters Slow Learning Rates from Kinematic Data

We observed that including kinematic data increased the accuracy in the static modality, but it came with the drawback of a greater training burden. However, it is not clear whether these effects are similar for the dynamic modality, in which the arm is dynamically moving, and inclusion of kinematic data could be more beneficial. Therefore, we trained classifiers on either the first static segment or the dynamic segment of dynamic modality, with including kinematic data, and tested on unseen data from both segments (figure 6(b)).

We found that, like the static modality, the gain in accuracy provided by the kinematic data inclusion is still present regardless of whether training was on the static or dynamic segments. All classifiers with kinematic data outperformed the models without the data, with a large number of trials for the training dataset. However, the increase in accuracy depended on which IMU was being included. The IMUs that increased the accuracy, as measured by the mean accuracy of both the models trained with the static and dynamic segments were trunk (93.2% ± 7.4%), arm (94.6% ± 6.4%), forearm (95.6% ± 5.3%), and wrist (96.7% ± 4.6%). Additionally, the models with multiple IMUs had even larger benefits (all except wrist: 98.0% ± 4.0%; and all: 98.9% ± 2.8%). These results are similar to those found in the static modality.

However, we found that training the models with different segments resulted in different learning rates. Using the τ parameter in equation (1), we found that the models without the kinematic data inclusion had an average τ value of 2.11 ± 0.72. However, the single IMU models trained with the static segment all had average values of at least 6.6, indicating slower learning rates (trunk: 7.3 ± 2.0; arm: 7.3 ± 0.9; forearm: 7.0 ± 0.8; and wrist: 6.6 ± 0.8). Notably, the trend changes with the single IMU models trained with the dynamic segment. These models have average values of around 5, a considerable increase in the learning rate (trunk: 5.3 ± 0.7; arm: 5.2 ± 0.6; forearm: 5.0 ± 0.6; and wrist: 4.7 ± 0.5; figure 6(b)). Therefore, the learning rates of the classifiers trained with the dynamic segment are faster than training with the static segments, when using classifiers that consider the kinematic data of the limb.

## 4. Discussion

By developing an open-source device to standardise data collection for myoelectric control, we confirmed the limb position effect from data collected from eighteen healthy subjects. The limb position effect in our data was similar to the prior research. Specifically comparing to [24], our data created a higher intra-position error (7.7.% compared to 3.8%) and lower inter-position error (18.4% compared to 21.2%). It is possible that this slight difference was due to the CAPSAS arm positions being two-dimensional versus the three-dimensionality of their experiment. However, overall, the limb position effect caused a significant increase in the errors of the classifiers in both studies.

To overcome the limb position effect while minimizing the training burden, we found that training on four different positions can be optimal. Indeed, prior research also had found that using data from multiple training positions can be beneficial [24-26]. Fougner et al. found a pattern of additional arm training positions decreasing the error, similar to what was shown in our work [24]. In particular, we found that four training positions are optimal for balancing the increased performance and the training burden drawback. Similarly, Fougner et al. found that increasing from three to five training positions decreased the error by only 0.4%.

Besides the limb position effect, which could be analysed exclusively using data from the static modality, we also examined the impact of the limb’s motion on myoelectric control using the data from our dynamic and object modalities. We showed that both static training in multiple positions and dynamic training could provide accurate control. Radmand et al. showed a similar result of dynamic training being as robust to the limb position effect as training in multiple positions [30]. However, their study was limited to one large dynamic motion. Our work used twelve different dynamic arm motions during the study, which allowed us to test the classifiers on a more extensive range of potential motions. Moreover, both their work and ours suggest that dynamic training is especially beneficial if the device is expected to be predominantly moving during control.

Current myoelectric controllers often map specific hand gestures to different commands in a prosthesis or an AR/VR environment. However, to create natural and intuitive interaction with the environment, the hand gestures performed should resemble the grasp required by the object’s shape. Our CAPSAS modular design allowed us to assess the myoelectric control accuracy while the participants were interacting with real-world objects. We found that the classifiers obtained during object modality had relatively poor accuracies compared to the traditional hand gestures used in most studies. The problem could arise from less distinct patterns appearing in the muscle activation since multiple sets of muscles are targeted (unlike the traditional hand gestures, which activate only a limited and specific set of muscles). Furthermore, since we did not dictate the exact fingers that should be involved during object grasps, it is possible that the individual differences and even within-session variability in grasping real-world objects impaired the classifier’s accuracy. Indeed, there are significant individual differences in grasping real objects such as a pen [44]. Such an issue could potentially be solved with more complex classification algorithms, although this solution would create new issues of power consumption and computational limits in embedded systems.

We next examined whether including the IMU data would enhance the classifier’s accuracy, and whether this inclusion would affect both static and dynamic modalities similarly. We found that including the IMU data in the classifier increased the accuracy. Similar patterns have also been documented by prior studies [24, 30, 31]. However, the increase in accuracy came with significantly worse learning rates, necessitating more training data and increasing the training burden. Comparing the different IMUs, we found that the wrist IMU will provide the most benefit to classification. However, the wrist IMU may not be possible in all myoelectric control applications, such as prosthesis control for individuals with higher-level amputation than wrist articulation. Therefore, we found that the next best option would be a forearm IMU, and then an arm IMU.

Moreover, we found that the learning rates of the classifiers appended with kinematic data can be improved with dynamic training. Earlier findings also back up these results. For instance, Radmand et al. hypothesised that dynamic training helped cover more positions for a classifier to learn faster [30]. From our study, we can conclude that this applies to classifiers that contain both myoelectric and kinematic information. We hypothesise that dynamic training allows the classifier to learn the position-specific muscle activation patterns in a faster manner than static training, although further research could be considered to solidify this hypothesis.

Consequently, based on our in-depth analysis, we recommend employing an inertial measurement unit (IMU) with dynamic training for optimal myoelectric control. In non-rehabilitation applications, wrist IMUs are particularly effective. Forearm IMUs also remain the most beneficial in a prosthesis application. The dynamic training approach is essential when incorporating IMU data for faster learning rates that will prevent an increased training burden, but will also enhance performance during activities with limb movement.

Some limitations of our study include the lack of individuals with amputations to generalize the findings readily to prosthesis control. If this study were solely focused on rehabilitation, it would be beneficial to include such participants. While not presented in this paper, others could replicate our developed open-source device to contribute to such a study. Future studies should also consider multiple modalities in a single session. This is indeed important because even though we used anatomical markers to place the EMG electrodes in the exact locations across days, electrode shift could still exist. Therefore, data collection for different modalities in the same session can reduce that confounding factor.

## 5. Conclusion

The effects of limb position and motion on myoelectric control were thoroughly studied. Classifiers had significantly decreased accuracy when tested in untrained positions, confirming the limb position effect. Depending on the context of the myoelectric control application, classifiers should be trained with four static positions or dynamic limb movements between those positions. Furthermore, appending kinematic data to the classifier increased its accuracy but reduced learning rates. An approach of training with dynamic data was shown to improve the learning rates of the classifier optimally. Hence, using dynamic training with kinematic data provides gains in accuracy while avoiding the most considerable learning rate impact.

Our open-source, automated CAPSAS device will help standardise datasets between labs, aiding the development of robust and widespread myoelectric control. The improved training strategies presented in this work could help lead the path towards creating more robust myoelectric control of devices, with applications ranging from more reliable prosthetic hands to control of mobile devices and AR/VR interactions.

## Acknowledgements

The authors declare that they have no competing interests related to this work. The authors would like to thank all the participants who took part in the study. Raelyn Tobillo is thanked for her contribution to the initial kinematic data collection. T.O. gratefully acknowledges the UCF SURF program for providing funding during this research. Some figures were created with assistance from Adobe Illustrator’s generative AI.

T.O. and M.R. conceived and planned the experiments.

T.O. and Z.A. created the data acquisition software. T.O., S.R., and J.W. collected the data. T.O. completed the data analysis with assistance from Z.A. All authors contributed to the manuscript. M.R. supervised the project.

