## Supplementary figures for "Limb Position Effect in Myoelectric Control: Strategies for Optimisation and Standardisation"

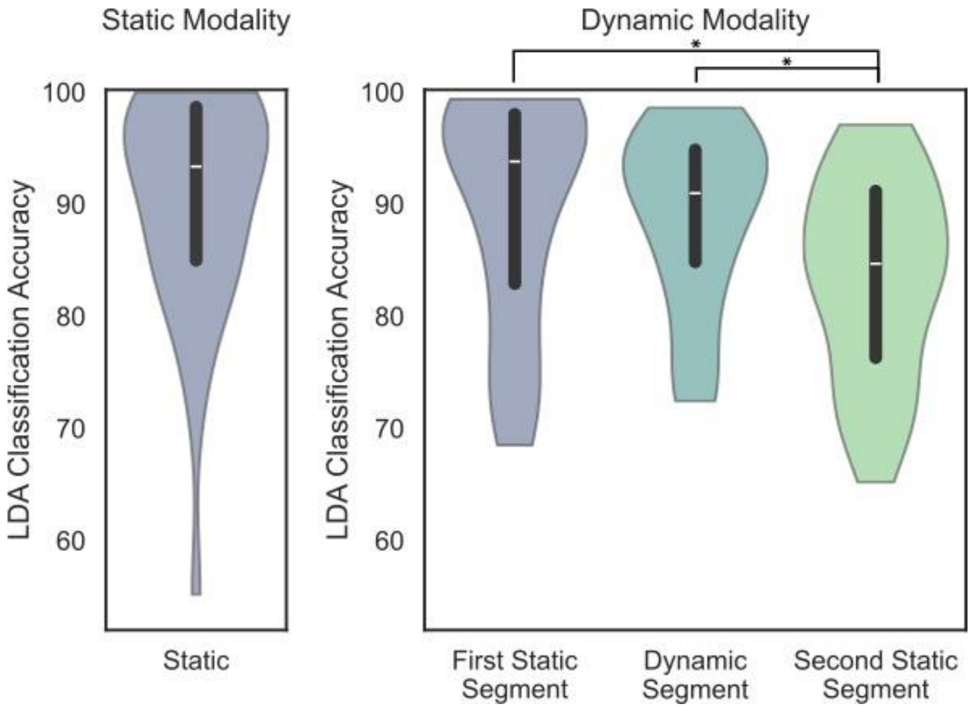


**Supplementary Figure 1.** Analysis of the segments of the dynamic modality compared to the static modality. For each modality, classifiers were trained on the outer four corners of the CAPSAS using a one-and-a-half-second window starting one second after the trial began (for the dynamic modality, this is the first static segment). The training dataset consisted of fifty trials and the testing dataset consisted of all remaining trials from that session. The static modality's testing dataset consisted of the remaining trials windowed using the same method as the training. The dynamic modality's testing datasets consisted of the remaining trials windowed in three parts based on the movement of the arm (first static segment, dynamic segment, and second static segment). To remove any motion artefacts from the two static segments, we only included a one-and-a-half-second window, which was one second after moving to the intended position and half a second before the end of the segment. Furthermore, to ensure the dynamic segment had motion, we excluded the first half-second of data from when CAPSAS signalled the participant to move. To ensure the number of data points between each static segment and the dynamic segment is equal, we also used a one-and-a-half-second window for the dynamic segment. Boxplots show the accuracy of the classifiers when tested under different conditions. Compared to the second static segment, the first static segment ($p = 2.0 \times 10^{-5}$, $r = 0.754$) and dynamic segment ($p = 4.6 \times 10^{-6}$, $r = 0.810$) perform significantly better (Wilcoxon signed-rank test).

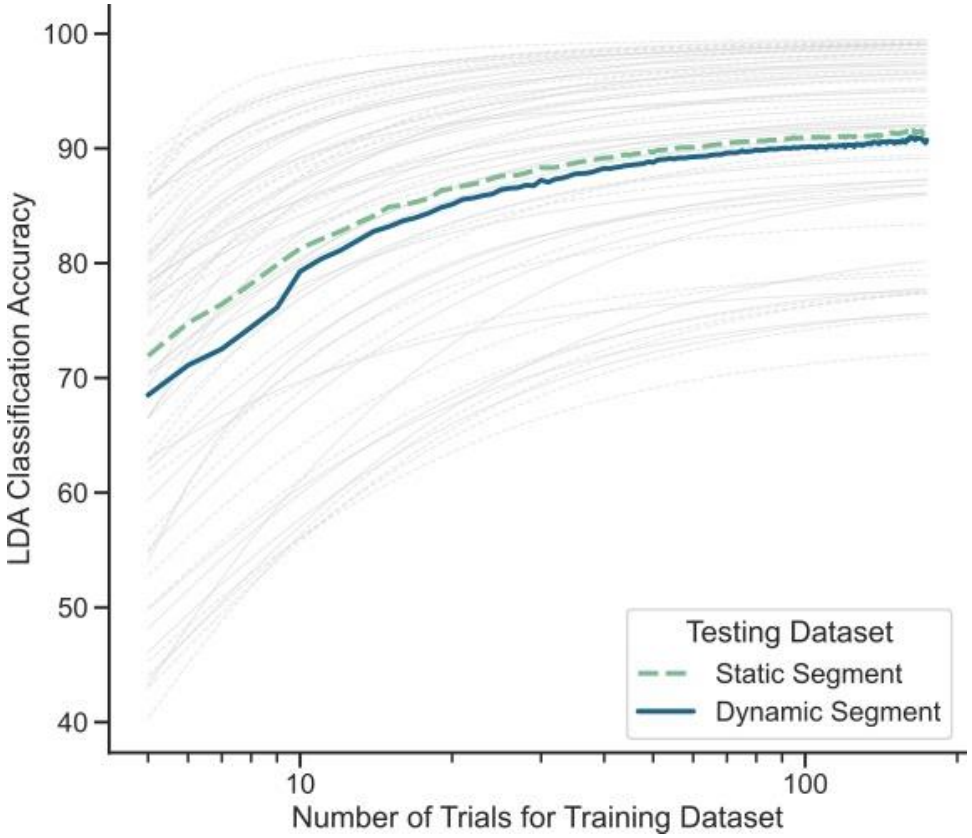


**Supplementary Figure 2.** Analysis of the effects of dynamic limb motion on classification accuracy. An LDA classifier is trained on only the first static segment of the dynamic session and tested on either the first static (dashed) or dynamic (solid) segments. The learning curves of individual sessions are fit to a hyperbolic curve (equation 1) and are drawn by thin grey lines (dashed or solid for static or dynamic, respectively). The mean learning curve of all participants for each test condition is coloured. The classifiers were tested on all remaining data.

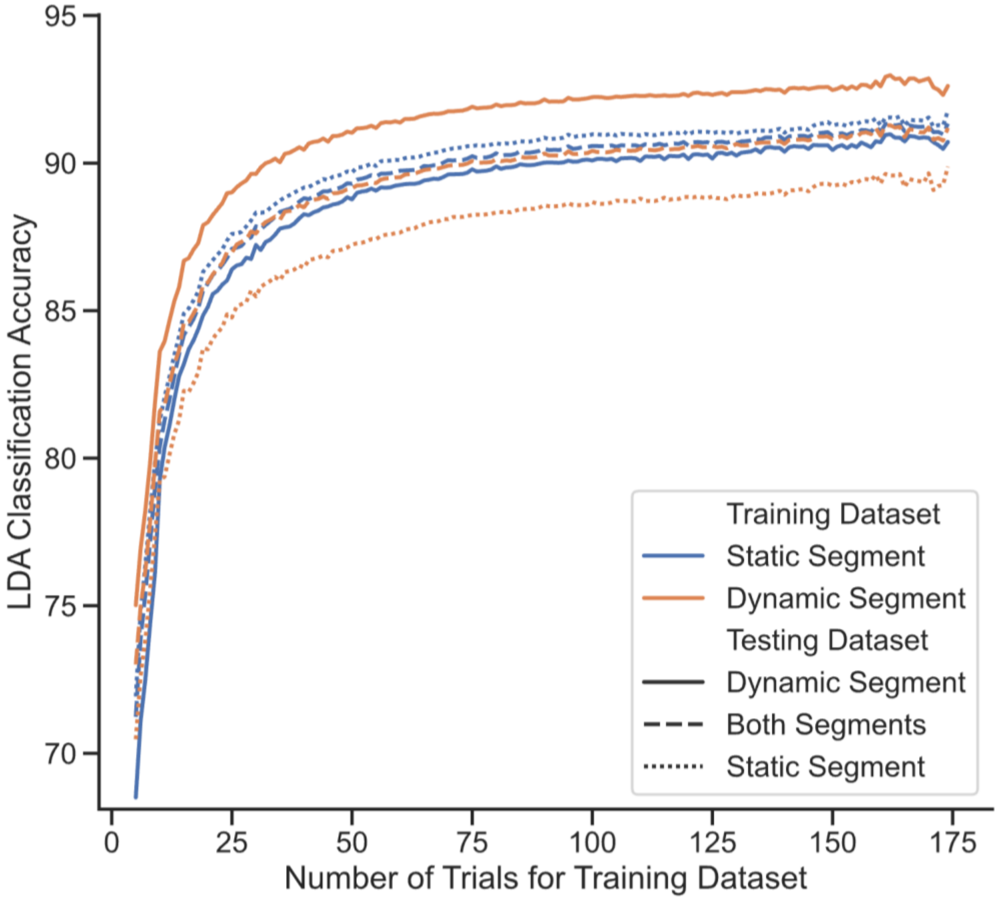


**Supplementary Figure 3.** Using data from the dynamic modality, a comparison of classifiers is made with specific training and testing segments. The mean classification accuracy for each training and testing type is shown as a line. Colour shows which segments of data are used for the training dataset. Solid lines show classifiers tested on only the dynamic segment, long dashed lines show classifiers tested on both segments, and short dashed lines show classifiers tested on only the static segment. The classifiers were tested on all remaining data.

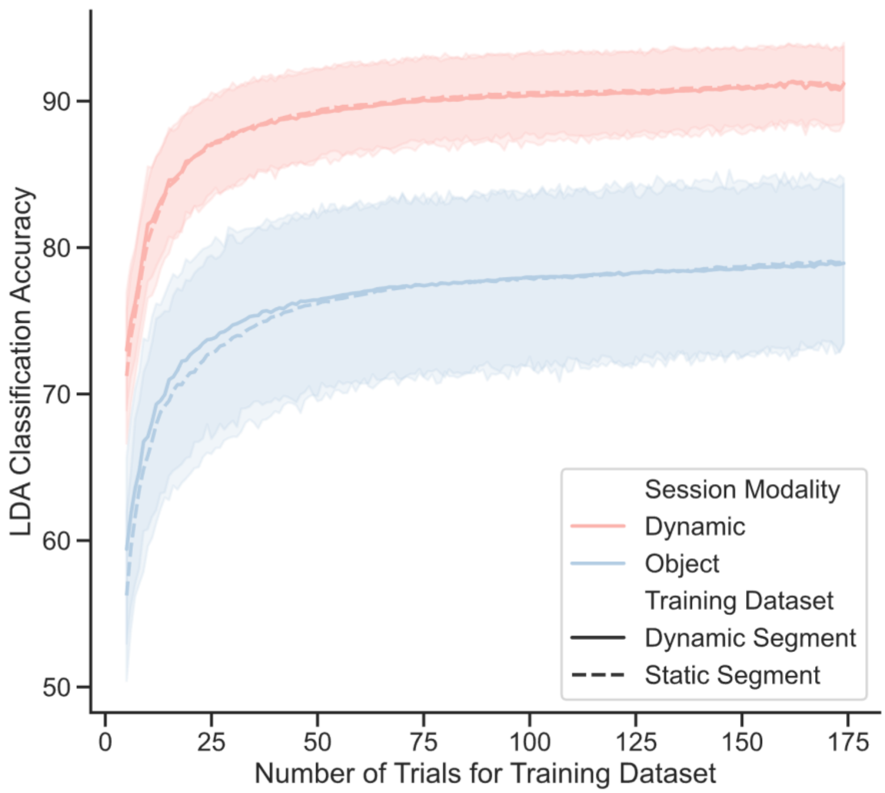


**Supplementary Figure 4.** Mean learning curves of the LDA classifiers are shown, with the dynamic modality outperforming the object modality. Dashed lines represent classifiers trained on the static segment of their corresponding modalities, while solid lines represent classifiers trained on the dynamic segment data. Colour represents which modality is being considered. The classifiers are tested with both segments from all the remaining trials. The shading shows a 95% confidence interval for each curve.
